# Dorsoventral gradient of theta sweeps in medial entorhinal cortex

**DOI:** 10.64898/2026.03.08.710351

**Authors:** Zilong Ji, Huiwen Zhang, Rokas Stonis, Neil Burgess

## Abstract

During random foraging, the positional signal decoded from entorhinal grid cells exhibits left–right theta sweeps, alternating from one side of the heading direction to the other across successive theta cycles. Here, we report that theta sweeps are topographically organised along the dorsoventral axis of the medial entorhinal cortex, with the angular deviation from heading direction increasing gradually from dorsal (smaller scale) to ventral (larger scale) modules. This gradient coexists with a corresponding dorsoventral increase in angular deviation decoded from theta-modulated directional cells, which drive grid-cell theta sweeps. These phenomena parallel a broadening of head direction tuning and increasing occurrence of theta cycle skipping in single cell firing along the dorsoventral axis. Computational modelling demonstrates that these patterns are consistent with continuous attractor dynamics and a dorsoventral gradient in firing rate adaptation. These results highlight how theta sweeps can simultaneously represent multiple potential future locations and reveal a clear neural mechanism underlying this process.

## Introduction

Theta sweeps are dynamic neural representations observed in the hippocampal formation during the theta rhythm (6-12 Hz). In place and grid cells, theta sweeps represent internally generated trajectories that extend outward from the animal’s current location. The trajectory in one theta cycle can appear as forward-directed progression in linear mazes [1], as left-right alternating progressions in T-shape mazes [2,3] and open-field environments [4]. They are driven by theta-modulated directional cells, which represent internal directional sweeps from side to side of the heading direction [4]. These population-level phenomena correspond to single-cell firing properties identified in earlier studies, including theta phase precession in place [5], grid [6] and theta-modulated directional cells [7], as well as theta skipping (i.e., firing on alternate theta cycles) in these cell types [3,7–11].

Theta sweeps reflect sequential activation of cell ensembles within individual theta cycles. This temporal organisation occurs on a similar time scale as spike-time-dependent plasticity [12], suggesting a mechanism by which theta sweeps could support temporal sequence learning and memory encoding [1,13–15]. Theta sweeps during open-field navigation may facilitate efficient sampling of space by transiently activating cells representing locations on the sweep path, thereby reducing the need for exhaustive behavioural exploration [4,16] and possibly aiding the formation of cognitive maps. Furthermore, recent findings indicate that theta sweeps in open-field navigation not only alternate stereotypically between left and right but can also encode directions toward future goal locations, independently from head or movement direction [17–19].

The hippocampal formation exhibits multiple forms of topographical organisation along the dorsoventral axis. In the medial entorhinal cortex (mEC), grid cells are arranged into discrete modules, with grid spacing increasing discontinuously from dorsal to ventral regions [20,21]. In the hippocampus, place cells show a corresponding dorsoventral increase in firing field size [22–24]. Similarly, in mEC layer III, head-direction cells exhibit sharper tuning dorsally and broader tuning ventrally [25]. In addition to these functional gradients in spatial tuning, intrinsic physiological properties are also spatially organised. For example, firing rate adaptation in mEC layer II principle cells increased progressively towards ventral regions [26]. Membrane conductance in layer II stellate neurons likewise exhibit a gradient, potentially reflecting gradients in HCN and leak potassium channels [27], as does the frequency of subthreshold membrane potential oscillations [28]. However, it remains unclear whether population-level dynamics are similarly structured along the dorsoventral axis, and which neural mechanisms might underlie such organisation. Our recent work identified firing rate adaptation as a potential mechanism generating theta sweeps at the population level [16], suggesting that circuit-level dynamics may also be topographically organised as a consequence of the graded organisation observed at the single-cell level. To address this question directly, we systematically examined whether population-level theta sweeps exhibit a structured gradient along the dorsoventral axis of the medial entorhinal cortex.

Here, we report that, in rats foraging for randomly scattered food, left–right theta sweeps in grid cells exhibit a pronounced gradient along the mEC dorsoventral axis, with sweep angles increasing from dorsal to ventral. A corresponding gradient is also found in theta-modulated direction cells (tmDCs) in the parasubiculum, where sweep angles similarly increase ventrally. These findings indicate that the geometric structure of theta sweeps varies systematically across the mEC and parasubicular dorsoventral axis, paralleling previously known gradients in single-cell features including grid spacing and head direction tuning [20,25], and suggesting that grid cell firing in different modules does not correspond to a common location at later phases of theta. We further suggest that this gradient in sweep angles may arise from a dorsoventral increase in neuronal firing rate adaptation [26], which has been proposed to generate theta sweeps in a ring-attractor model of tmDCs and to drive corresponding theta sweeps in downstream grid cells [16]. Such organisation may enable the brain to simultaneously represent multiple future potential locations in the environment than if dorsoventral theta-sweep directions aligned.

## Results

### Topography of sweep angles in grid cells

We analysed data from Vollan et al. [4], comprising 29 open-field running sessions recorded from 17 Long–Evans rats. To decode theta sweeps, we followed the method described by Vollan et al. [4], correlating the instantaneous population activity (PV) of all recorded cells in each temporal bin with the session-averaged population activity in each spatial bin (see Methods). The decoded position was then assigned to the spatial bin showing the highest correlation score across all bins. Using this approach, we replicated the decoding of left-right theta sweeps: across successive theta cycles, the decoded position swept outward from the animal’s location into the surrounding environment, with the sweep direction alternating between left-forward and right-forward (Fig. 1A&B). The averaged sweep angles range from 5.0° to 47.8° across all recording sessions (median = 24.1°; IQR: [16.1°, 29.0°]), like results reported in Vollan et al. [4].

**Fig. 1:**
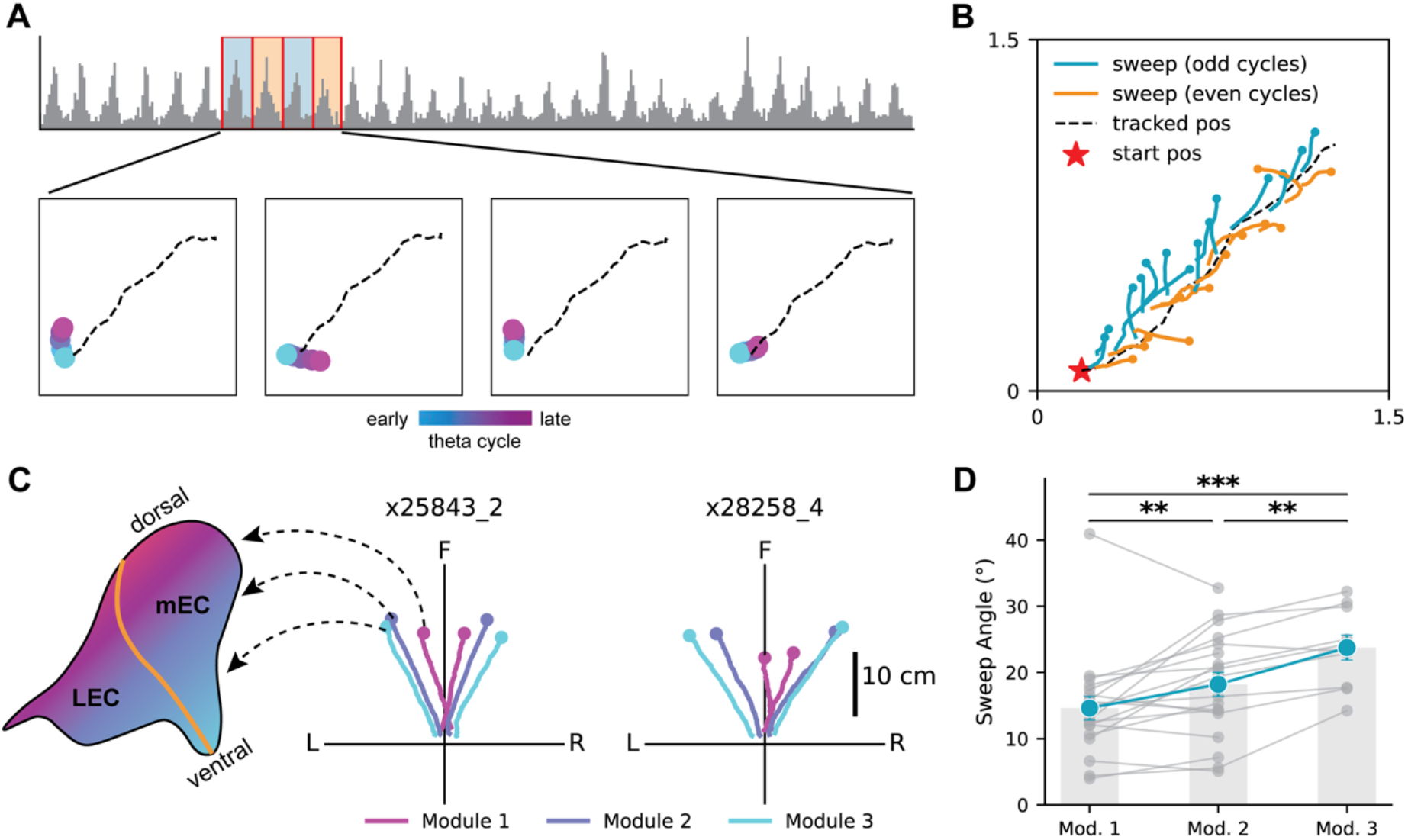
Topography of sweep angles in grid cells along the mEC dorsoventral axis. **(A)** Example of left– right alternation across four successive theta cycles, reproduced from Vollan et al [4]. Top: summed spike counts of all recorded cells over a short period, displaying theta rhythmicity. Bottom: decoded sweeps in four consecutive theta cycles, with dashed lines indicating the animal’s trajectory over the same period as in the top plot. **(B)** Decoded sweeps across the recording period showing in (a), with blue denoting left sweeps and orange denoting right sweeps. **(C)** Left: a schematic outline of the left entorhinal cortex in a rat brain. The dorsoventral gradient of the entorhinal cortex (magenta to blue) spans both the lateral and medial entorhinal subdivisions (lEC and mEC), with the border indicated by the orange line. Right: Mean left–right sweeps decoded from grid module 1 (magenta), module 2 (purple) and module 3 (blue), in two animals. **(D)** Mean sweep angle across three grid modules. Each dot represents the sweep angle decoded from one grid module (48 modules in total). Grey lines link the mean sweep angles from grid modules in the same recording session (19 sessions from 11 animals). A linear mixed-effects model yielded a significant effect of module identity on sweep angle, with *χ*^2^(2) = 22.7, p = 1.2 × 10^−5^. Averaged sweep angles were module 1, 14.6°; module 2, 18.2°; module 3, 23.8°. Module 2 vs module 1: β = 3.6, p = 0.003. Module 3 vs module 2: β = 5.0, p = 0.001.

Since grid cells are organised into multiple grid modules along the mEC dorsoventral axis [20,21], we next asked whether sweep features are different across grid modules. To do this, we decoded theta sweeps from grid cells in each module independently. Only sessions containing at least two grid modules, each with no fewer than 40 grid cells, were included in the analysis (19 sessions from 11 animals, 48 modules in total). This resulted in 42 to 215 grid cells per module (median = 100; IQR: [69-129]). We then calculated the mean sweep angle for each module and found that the sweep angle increased across grid modules along the dorsoventral axis (see Fig. 1C for results from two rats). A linear mixed-effects model (LMM) revealed a significant effect of module identity on sweep angle (Fig.1D; χ^2^(2) = 22.7, p = 1.2 × 10^−5^. Averaged sweep angles were module 1, 14.6°; module 2, 18.2°; module 3, 23.8°. Module 2 vs module 1: β = 3.6, p = 0.003. Module 3 vs module 2: β = 5.0, p = 0.001.).

To control for potential confounds arising from unequal cell numbers across modules— which could bias the decoded sweep angle (fewer cells leading to noisier decoding)—we subsampled each module to match the number of grid cells in decoding and repeated the analysis. The results still showed a significant module effect and revealed a systematic increase of sweep angles along the mEC dorsoventral axis (LMM with (*χ*^2^(2) = 14.7, p = 6.4 × 10^−4^; Fig. S1). Furthermore, a recent study reported that sweep angle varies with theta frequency during REM sleep [19], with larger sweep angles observed during slow theta oscillations (< 6.5 Hz) than during fast theta oscillations (> 8.5 Hz). We therefore examined the theta frequency derived from the summed spike activity in each module and did not observe any significant differences between modules (Fig. S2; LMM with χ^2^(2) = 4.77, p = 0.09). These results ruled out a possibility that a gradient decrease in theta frequency along the dorsoventral axis accounts for the increase in sweep angle observed in the data.

### Grid-cell sweep-angle topography reflects that of theta-modulated directional cells

Sweeps in grid cells align with those in theta-modulated directional cells (tmDCs; 20 to 450 per recording session, median = 194; IQR:[85-287]; Fig. 2A) [4], which have been identified in the anteroventral thalamus nuclei [11,29], parasubiculum and mEC [4,10,30]. First, we showed that the decoded direction (hereafter referred to as the internal direction) sweeps smoothly around the head axis (Fig. S3), rather than flickering abruptly from side to side of the head axis [4], across successive theta cycles. Furthermore, it has been shown that the locational sweep direction decoded from grid cells matches the internal direction decoded from tmDCs (Fig. 2B) [4]. We next asked whether tmDCs exhibit a gradient in sweep angle along the dorsoventral axis.

**Fig. 2:**
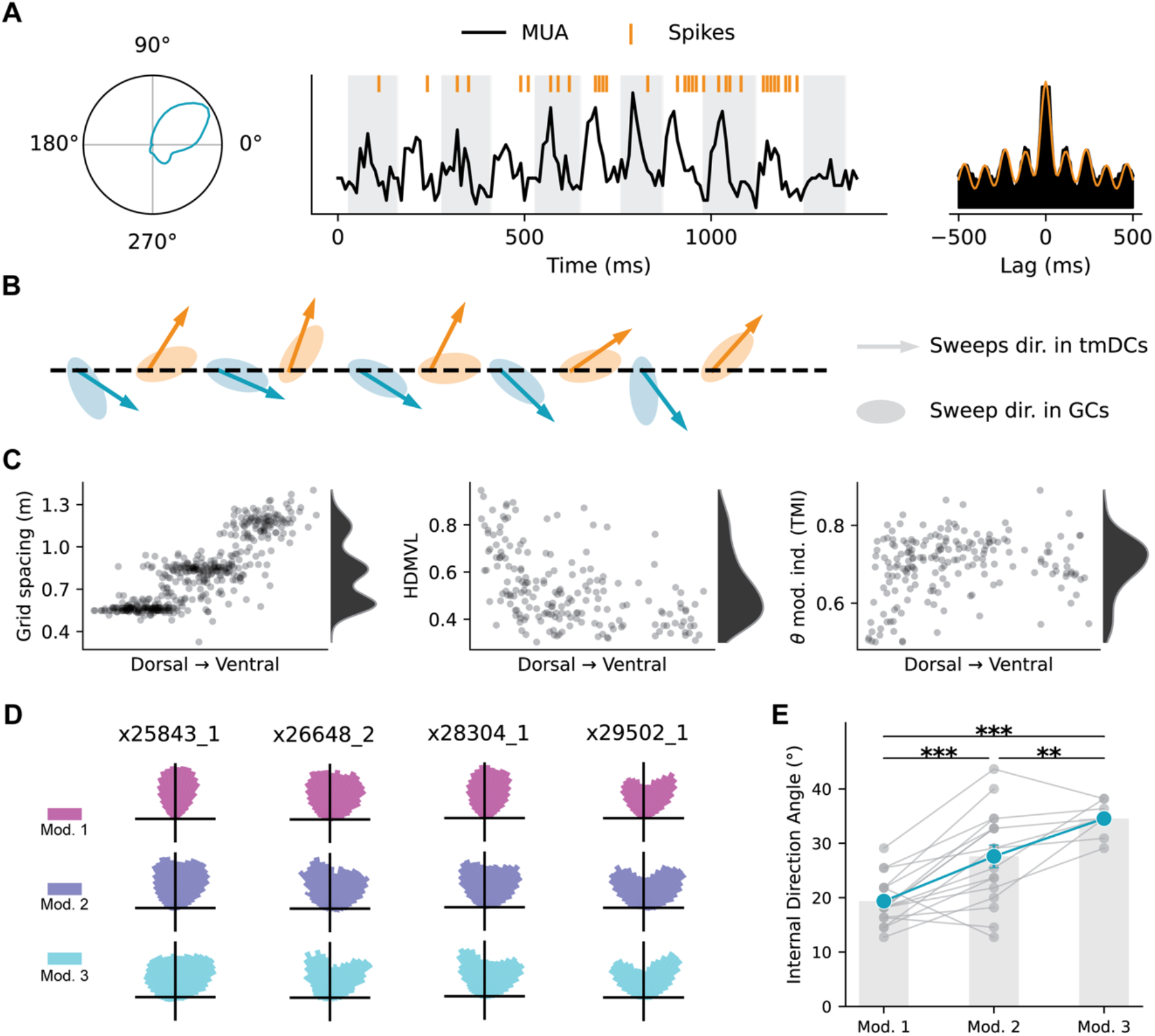
Topography of sweep angles in tmDCs along the parasubicular-mEC dorsoventral axis. **(A)** Left: a theta-modulated direction cell with its directional tuning field relative to head direction. Middle: a segment of the cell’s spike train overlaid on summed spike counts of all co-recorded cells with theta cycles marked by alternating white and grey boxes. Right: the autocorrelogram of the cell’s spike train from the full recording session, with the orange line a cosine wave fit. **(B)** Example epochs of decoded sweep direction in tmDCs (arrows) and in grid cells (patches) shown across 13 successive theta cycles, reproduced from Vollan et al [4]. **(C)** Left: distribution of grid spacing at different recording sites along the dorsoventral axis from one recording session, with each dot representing a grid cell. Middle: distribution of head-direction mean vector length (MVL) calculated for each tmDC from the same session. Right: distribution of the theta modulation index calculated for each tmDC from the same session. **(D)** Polar histograms of decoded sweep angles in tmDCs for each assigned module, with each row corresponding to one module and each column to data from one animal. **(E)** Mean sweep angle across the three assigned modules. Each dot represents the sweep angle decoded from one recording session (17 sessions from 10 animals). Grey lines link the mean sweep angles from assigned modules in the same recording session. A linear mixed-effects model yielded *χ*^2^(2) = 30.7, p =2.1 × 10^−7^. Averaged sweep angles were module 1, 19.4°; module 2, 27.6°; module 3: 34.5°. Module 2 vs module 1: β = 8.2, p = 1.5 × 10^−6^. Module 3 vs module 2: β = 6.2, p = 0.006.

First, we examined whether tmDCs are organised into modules, as reported for grid cells [20]. We used two criteria to assess this effect: the head-direction mean vector length (HDMVL) and theta modulated index (TMI; see Methods). Neither of these measurements show any discontinuity along the dorsoventral axis, unlike grid scale (Fig. 2C for an example from one animal and Fig. S4 for more animals). Therefore, to perform population decoding of tmDCs at different locations along the dorsoventral axis, we assigned them into different “modules” according to their nearby grid cells. Specifically, for each recording, we trained a logistic regressor to predict the grid-module identity of each grid cell from its dorsoventral recording position (see Methods). Each tmDC was then assigned to a grid module by predicting the module identity from its recording position. This results in 2 to 251 tmDCs in each assigned module (median = 63; IQR: [36-107]). At least two assigned modules per session were required for within-animal comparison of directional sweep angles.

Next, we decoded the internal direction from the population activity of tmDCs within each assigned module. Modules were included only if they contained at least 20 tmDCs, yielding 42 modules across 17 sessions from 10 animals (see Methods). To compute a single directional sweep angle for each module, we identified the theta phase at which tmDC population activity peaked, extracted the corresponding internal directions across theta cycles, and computed their circular mean. We found a systematic increase in the directional sweep angle across assigned modules along the dorsoventral axis (Fig. 2D&E; A linear mixed-effects model (LMM) with *χ*^2^(2) = 30.7, p = 2.1 × 10^−7^, internal-direction sweep angles were module 1, 19.4°; module 2, 27.6°; module 3, 34.5°. Module 2 vs module 1: β = 8.2, p = 1.5 × 10^−6^. Module 3 vs module 2: β = 6.2, p = 0.006). To control for potential differences in cell numbers, we subsampled each module to match the number of tmDCs used for decoding, and the results still showed a significant modular effect and revealed a systematic increase of sweep angles along the parasubicular/mEC dorsoventral axis (LMM with *χ*^2^(2) = 27.5, p = 1.1 × 10^−6^; Fig. S1).

For the analysis above, the same theta-phase window was used to extract directional sweep angle across all assigned modules, which corresponds to the centre (180° phase) of each LFP theta cycle. However, we also observed that tmDCs within each assigned module fired at slightly different phases of the global reference theta cycle––dorsal cells at earlier phases and ventral cells at later phases (Fig. S5)––in line with the gradual theta-phase shift along the mEC dorsoventral axis [31]. To account for this variation, we extracted the directional sweep angle for each assigned module using the phase window corresponding to its own peak activity within the theta cycle. The dorsoventral gradient in directional sweep angles remained (LMM with *χ*^2^(2) = 25.3, p = 3.2 × 10^−6^; Fig. S5).

### Topography of directional sweeps reflects topography of single-cell firing features in tmDCs

The topography of directional sweeps in tmDCs mirrors corresponding gradients in single-cell firing properties along the dorsoventral axis. First, due to the left-right alternation of directional theta sweeps [4], a tmDC can be activated even when the animal’s head is not pointing towards its preferred firing direction, which broadens the head-direction tuning field and reduces the mean vector length (Fig. 3A). As sweep angles are larger ventrally, this reduction should be stronger in ventral tmDCs. Consistent with this prediction, we observed a progressive decrease in head-direction mean vector length (HDMVL) across assigned modules along the dorsoventral axis (Fig. 3B; see also Fig. 2C; A linear mixed-effects model (LMM) yielded *χ*^2^(2) = 415.0, p =7.6 × 10^−91^; HDMVL values were module 1, 0.574; module 2, 0.482; module 3, 0.461). This provides a mechanistic explanation for the reported decrease in HDMVL along the dorsoventral axis [25]. Apart from assessing head-direction tuning (plotting firing rate against the head direction), we also examined internal-direction tuning (plotting firing rate against the internal direction from the assigned module the tmDC belongs to) (Fig. 3A). Interestingly, the internal-direction mean vector length (IDMVL) did not show a gradient (Fig. 3C; LMM with *χ*^2^(2) = 2.60, p =0.27). This supports the idea that tmDC activity is driven more strongly by internal direction than by heading direction, and that the broadening in head-direction tuning reflects the increasing difference between the two.

**Fig. 3:**
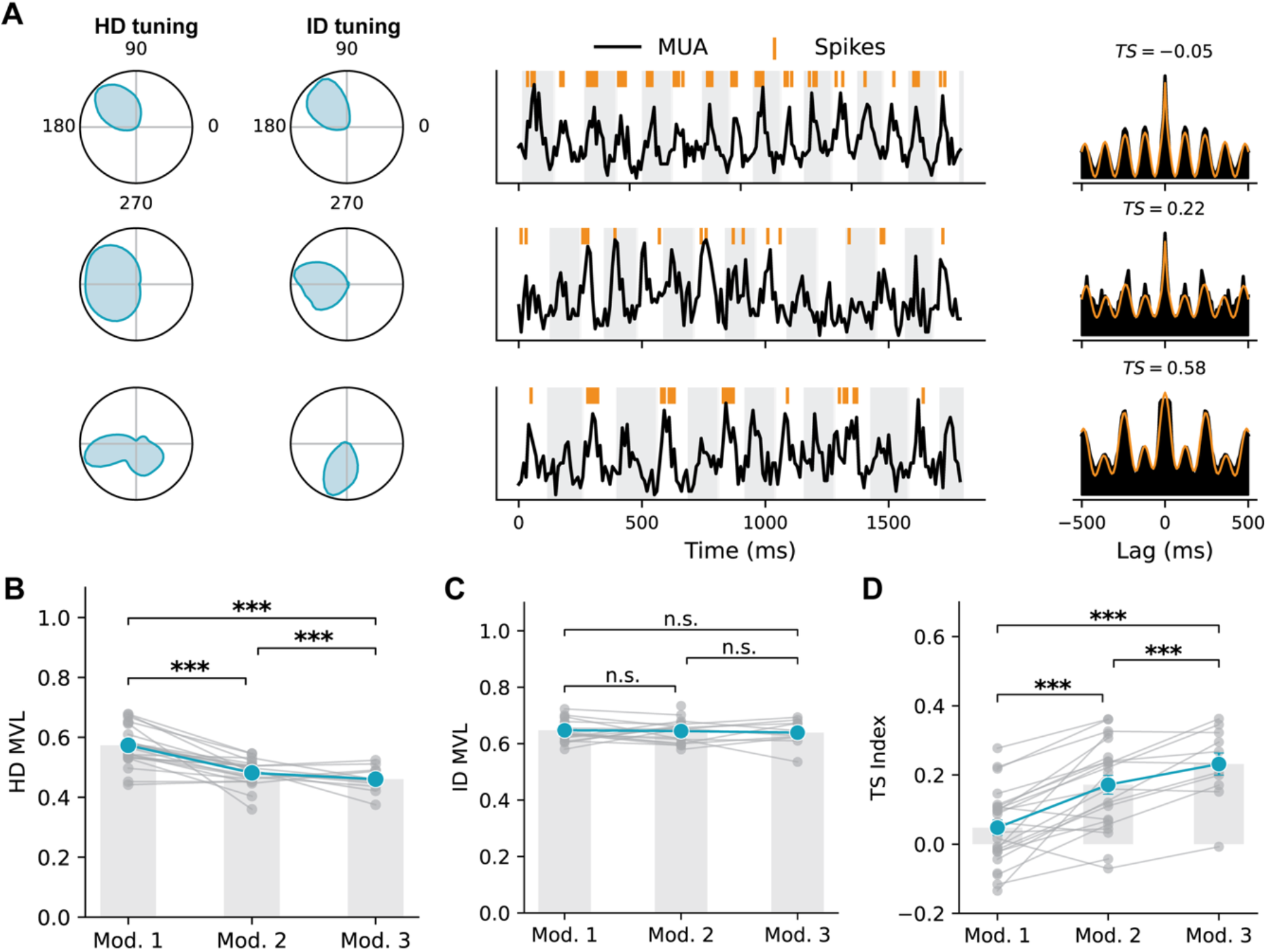
Topography of single-cell firing features in tmDCs along the parasubicular-mEC dorsoventral axis. **(A)** Three example tmDCs from each of the three assigned modules in the same recording session. Column 1: directional tuning fields relative to the head direction. Column 2: directional tuning fields relative to the decoded internal direction. Column 3: segments of cells’ spike trains overlaid on the local field potential, with theta cycles marked by alternating white and grey boxes. Column 4: autocorrelograms of the cells’ spike trains from the full recording session, with each orange line a cosine wave fit. **(B)** Head-direction mean vector length (HDMVL) across assigned modules. Each dot represents the mean HDMVL from all tmDCs in one assigned module (42 modules from 17 sessions). Grey lines link the mean HDMVL values from assigned modules in the same recording session. A linear mixed-effects model (LMM) yielded *χ*^2^ (2) = 432.6, p = 1.2 × 10^−94^. Mean HDMVL values were module 1, 0.574; module 2, 0.482; module 3, 0.461. Module 2 vs module 1: β = 0.086, p = 3.35 × 10^−63^. Module 3 vs module 2: β = 0.018, p = 8.87.6 × 10^−4^. **(C)** Internal-direction MVL (IDMVL) across assigned modules (LMM with *χ*^2^(2) = 2.60, p = 0.27). **(D)** theta skipping index (TSI) across assigned modules. A LMM yielded *χ*^2^(2) = 629.3, p = 2.2 × 10^−137^. Mean TSI values were module 1, 0.048; module 2, 0.172; module 3, 0.232. Module 2 vs module 1: β = 0.097, p = 4.89 × 10^−54^. Module 3 vs module 2: β = 0.071, p = 3.52 × 10^−27^.

Second, because theta sweeps in tmDCs alternate from side to side of the heading direction across successive theta cycles, especially during straight runs [4,16], this predicts that a tmDC with its preferred firing direction on one side of the current heading direction should be activated on every other theta cycle—a phenomenon known as theta skipping (Fig. 3A) [8–11]. As sweep angles are larger ventrally, tmDCs in the ventral part should display a stronger theta skipping effect. Consistent with this prediction, we observed an increase in the theta skipping index (TS index; see Methods) across assigned modules along the dorsoventral axis (Fig. 3D; LMM with *χ*^2^(2) = 629.3, p =2.2 × 10^−137^, mean TS index values were module 1, 0.048; module 2, 0.172; module 3, 0.232). Together, these single-cell properties converge to demonstrate a dorsoventral organisation of directional theta sweeps in tmDCs.

### Dorsoventral increase in firing-rate adaptation can explain the gradient of theta sweeps in a coupled continuous attractor model

Left–right sweeps in grid cells can be modelled using a two-layer continuous attractor network (CAN; Fig. 4A) [16], following the circuit described by Vollan et al. [4]. The model comprises a ring attractor for tmDCs and a two-dimensional (2D) CAN for grid cells (see Methods). Introducing firing-rate adaptation (FRA) into the model causes the activity bump in the tmDC ring attractor—representing the animal’s internal estimate of heading—to oscillate around the true heading direction rather than remaining stationary. This behaviour arises from a dynamic competition between two opposing forces: sensory input, which anchors and stabilises the bump at the current heading direction, and firing-rate adaptation, which acts as a form of activity-dependent negative feedback. As neurons within the bump remain active, adaptation progressively reduces their firing, weakening the stability of the bump and pushing it away from its current position. The resulting interplay between stabilisation and destabilisation leads to a continual sweeping motion around the heading direction (Fig. 4C&G). This, in turn, drives the activity bump in the 2D CAN to move outward from the current position along the same internal direction, alternating between left and right (Fig. 4C&H).

**Fig. 4:**
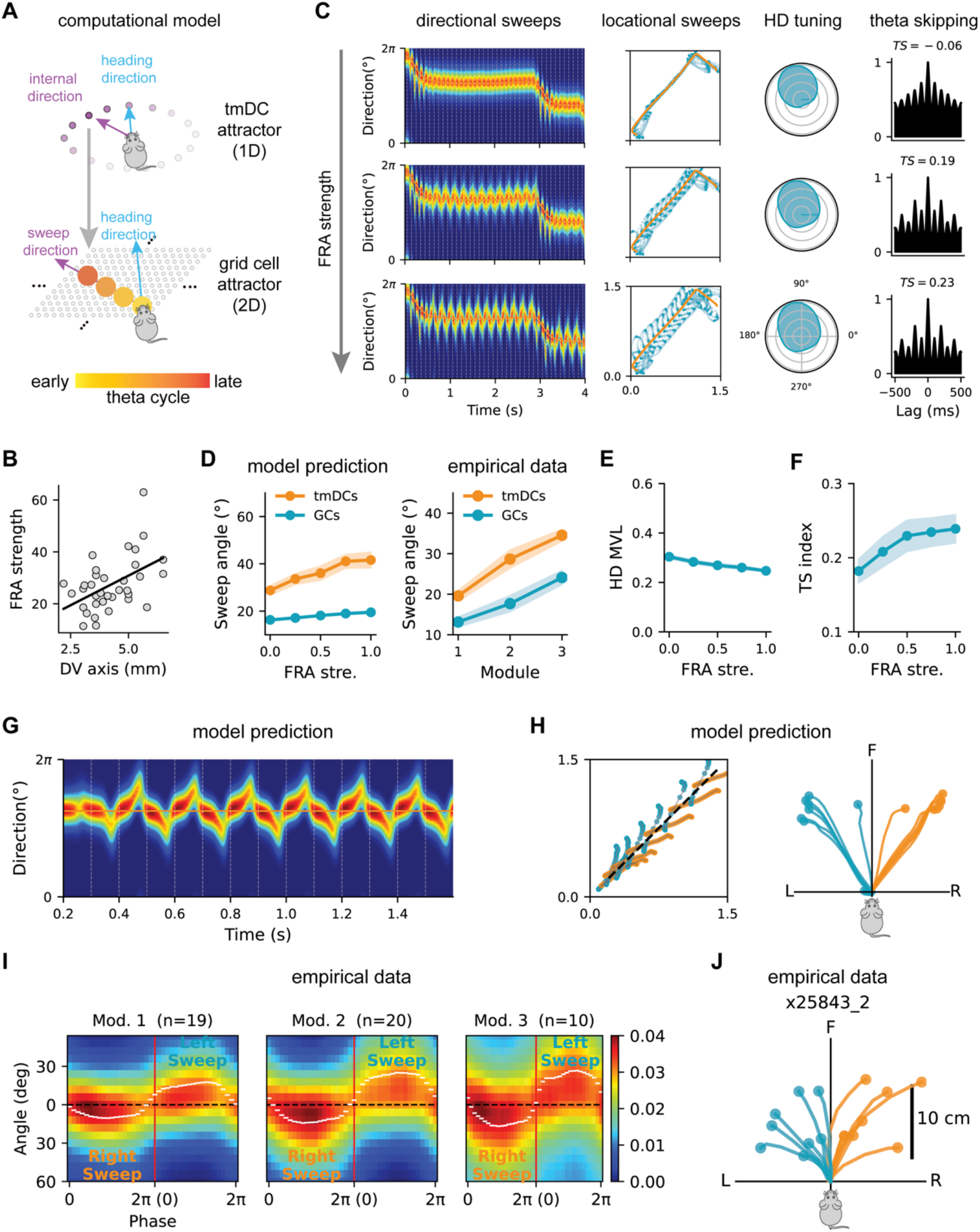
Dorsoventral increase in firing-rate adaptation (FRA) strength explains the gradient of theta sweeps and single-cell firing features in a coupled continuous attractor model of tmDCs and grid cells. **(A)** Schematic of the computational model, comprising a ring attractor of tmDCs (top) and a two-dimensional (2D) attractor of grid cells (bottom). The blue arrow marks the head direction, and the purple arrow marks the internal direction or sweep direction in a theta cycle. Circles from light to dark orange represent network activity in a theta cycle from early to late theta phases, sweeping in the same direction as the internal direction. **(B)** FRA strength increases with recording location along the dorsoventral axis for 35 mEC cells recorded using in vitro whole-cell patch-clamp electrophysiology (Pearson correlation: r = 0.48, p = 0.003). Data from Yoshida *et al*. [26] with permission. **(C)** Three modelling examples in which only FRA strength was varied. Column 1: directional sweeps in the tmDC ring attractor, with the orange line marking heading direction. Column 2: locational sweeps in the 2D grid-cell attractor (blue), with a segment of the simulated trajectory marked in orange. Column 3: firing field of a tmDC relative to head direction. Column 4: auto-correlogram of the cell’s firing. **(D)** Left: a model prediction of directional (orange) and locational (blue) sweep angles as a function of normalized FRA strength; right: directional (orange) and locational (blue) sweep angles as a function of the module index in empirical data. **(E)** Mean head-direction mean vector length across all simulated tmDCs as a function of FRA strength. **(F)** Mean theta-skipping index as a function of FRA strength. **(G)** Directional sweeps across theta cycles with a straight run in the model. **(H)** A model prediction of palm-tree-like sweep patterns in grid cells, overlaid on the straight trajectory (left) and reoriented with the heading direction pointing upwards (right). Backward sweeps were excluded. **(I)** Averaged directional sweeps in each assigned module in empirical data (merged across animals, with the module number indicated in brackets). The red line separates left and right directional sweeps. The dark dashed line marks the heading direction (reoriented to zero), and the white dashed line marks the mean directional sweep angle at each theta phase. **(J)** An example of palm-tree like locational sweeps decoded from grid cells in empirical data from animal x25843_2 and gird module 2. Sweeps were translated so that the starting point of each sweep was always at the origin and heading directions were re-oriented to the north.

In vitro whole-cell patch-clamp recordings [26] showed that FRA strength in principal neurons in mEC increases along the dorsoventral axis, measured as the ratio of the instantaneous firing rate between the first two induced spikes and the last two induced spikes (Fig. 4B). In the model, increasing FRA strength caused the activity bump in the ring attractor to sweep further away from the heading direction (Fig. 4C), thereby increasing the sweep angle in both the tmDC ring attractor and the grid-cell 2D CAN (Fig. 4D). Furthermore, since sweeps in grid cells are driven by upstream tmDCs, the model predicts larger sweep angles in tmDCs than in grid cells, which we verified empirically (Fig. 4D).

The network model also explains single-cell firing features in tmDCs we observed empirically. Specifically, increasing FRA strength increased the directional sweep angle, and therefore broadened the head-direction tuning curves of tmDCs and therefore reduced their head-direction mean vector length values (Fig. 4C&E). It likewise increased the theta-skipping index, reflecting the stronger influence of left-right directional sweeps under higher FRA strength (Fig. 4C&F).

Finally, since the tmDC ring attractor is a 1-dimensional continuous attractor network, we predicted that the activity bump should sweep continuously, rather than flickering between left and right (Fig. 4G). This prediction was verified empirically (Fig. 4I & Fig. S3). Interestingly, because the internal direction gradually deviates from the heading direction from the early to the middle phase in each theta cycle, we further predicted a gradual increase in the angle of locational sweeps across the early-to-middle phases of the theta cycle, resulting in a palm-tree-like pattern rather than straight locational sweeps (Fig. 4H). This prediction was likewise confirmed empirically (Fig. 4J & Fig. S3). In summary, this computational model provides a coherent mechanistic explanation for the empirical findings reported above and generates experimentally testable predictions.

## Discussion

This study provides empirical evidence for a topographical organisation of theta sweep angles and a potential mechanism underlying this organisation. Although topographical gradients in single cell tuning in mEC have been reported, e.g., grid spacing [20,21] and directional tuning breadth [25], gradients in the population representation of space have not. Using a large-scale dataset [4] together with computational modelling, we showed that: (1) theta-sweep angle increases dorsoventrally across mEC grid modules, thereby allowing different modules to represent multiple potential future locations simultaneously at late phases of theta; (2) this increase coexists with a corresponding dorsoventral increase in directional sweep angles decoded from tmDCs; (3) the gradient of theta sweeps accounts for the decrease in directional tuning and the increase in theta-cycle-skipping effect in tmDCs; (4) continuous directional sweeps drive locational sweeps, which exhibits a palm-tree like pattern; and (5) these features can be unified within an attractor model in which firing-rate adaptation increases along the mEC dorsoventral axis.

The computational model provides mechanistic explanations for empirical results we observed, as well as several experimentally testable predictions. First, it suggested that alternating theta sweeps in tmDCs and theta skipping are two manifestations of the same underlying mechanism. Since theta skipping was also observed in tmDCs in the anteroventral thalamus nuclei (AVN) [11,29], it raises the possibility that alternating directional theta sweeps originate in a region upstream of parasubiculum. It further predicted that when animals transition from straight running to turning, alternating directional sweeps should transform into forward-directed sweeps: the decoded direction should begin behind the current heading direction and propagate towards the future turning direction in each theta cycle [7], analogous to forward-directed theta sweeps observed in place cells during linear-track navigation [1]. This mechanism produces theta phase precession of turning angle during rotation in single-cell firing of tmDCs, observed recently in AVN [7]. Given the stronger alter-nating-sweep effect in more ventral modules, we would therefore expect the length of forward-directed theta sweeps during turning to also increase dorsoventrally. This, in turn, predicts a topographical organisation of theta phase precession in tmDCs during turning. Testing these predictions in tmDCs will be important when the full dataset from Vollan *et al*. [4] becomes available.

We propose that the topographical organisation of sweep angles reflects a gradient of firing-rate adaptation (FRA) strength along the dorsoventral axis in mEC [26]. This gradient likely reflects changes in biophysical properties of cell firing, specifically a decrease in the amplitude and an increase in the duration of the medium afterhyperpolarisation potential (mAHP) along the dorsoventral axis [26]. Several questions remain open. First, it is not yet clear whether the principal cells recorded in this region were grid cells. Second, little is known about FRA organisation in mEC layer III or the parasubiculum, which contain tmDCs [32]. Third, if the FRA hypothesis is correct, the underlying neurophysiological mechanisms that support its gradient along the mEC or parasubicular dorsoventral axis remain to be identified [27,33]. Addressing these issues will be essential for developing a detailed mechanistic understanding of the topographical organisation of theta sweep angles, and of the related single-cell firing features reported here.

Although theta sweeps can reduce the need for extensive behavioural sampling (e.g., navigating between locations) by virtually traversing space, sweeps with a fixed angle and distance would still cover only a limited portion of the environment. The present observation of simultaneous multi-angle theta sweeps—together with recent evidence that sweep length increases along the dorsoventral axis of the mEC [4]—suggests a more efficient sampling strategy. Dorsal sweeps are short and predominantly forward directed, providing fine-grained sampling ahead of the animal, whereas ventral sweeps are longer and extend laterally, capturing coarser information to the sides. Operating in parallel, these sweeps could link environmental features across multiple spatial scales and directions, enabling a cognitive map to form more rapidly than would be possible through physical exploration alone. This constitutes a working model for how multi-scale theta-sweep dynamics may facilitate rapid map formation. Finally, in addition to efficient sampling via stereotypic left–right alternating theta sweeps, theta sweeps in open-field environments can be goal-directed; that is, they can propagate from the animal’s position towards future goal locations, thereby providing a neural substrate for online goal-directed spatial planning [17–19].

## Materials and Methods

### Identifying theta-modulated direction cells

For calculating theta modulation index (TMI) for each cell, theta phase was extracted from the population spiking activity of all units. Specifically, spike times were binned into 10-ms bins and the resulting spike counts were bandpass-filtered between 5–10 Hz. PCA was applied to the matrix of bandpass-filtered spike counts with the first two principal components extracted. Theta phase at time *t* was then defined as the direction of the projection of the instantaneous population activity (PV) at time *t* onto the plane defined by PC1 and PC2. The phase of peak activity is roughly 180° relative to a theta cycle. For each neuron, a theta-phase tuning curve was constructed by calculating the spike rate in each theta phase bin and TMI was then quantified as the mean vector length (MVL) of the theta-phase tuning curve as:

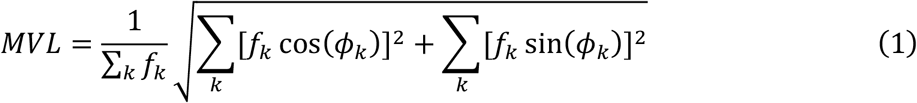

Where *f*_*k*_ is the firing rate in the *k*^*th*^ theta phase bin *ϕ*_*k*_.

For calculating head-direction mean Rayleigh vector length (HDMVL), the directional tuning curve relative to head direction was first computed by calculating the spike rate in 60 angular bins of head direction. Then HDMVL was calculated as the mean Rayleigh vector length using the firing rate in each head direction angular bin.

In our analysis, tmDCs are identified with two conditions: (1) theta modulation index (TMI) > 0.5; (2) HDMVL > 0.3. These two values are taken from the processed data in Vollan et al [4]. With these two criteria, we found 27 to 450 tmDCs across 24 sessions in 15 rats (median = 190; IQR: [96-235]).

### Determining modular identity for individual tmDCs

While tmDCs do not show modular organisation comparable to grid cells (Fig. 2C), we still wanted to decode directional representation from tmDCs at different recording locations along the dorsoventral axis. Therefore, we assigned module identities to each of the tmDCs by using their nearby grid cells which have been classified into grid modules in the Vollan data. Specifically, for each recording session, we trained a multinomial logistic regression model using the recording shank positions of grid cells as the predictor and their known module identities as the categorical outcome. We then applied the trained model to shank positions of tmDCs to assign their module identities. This resulted in 2 to 251 tmDCs in assigned modules (median = 63; IQR: [36-107]). Note that the fourth module, when present, was excluded from all analyses, as most recording sessions contained fewer than four grid modules.

### Decoding directional theta sweeps in tmDCs

To decode the sweep angle from tmDCs, we correlated the population activity vector (PV) from selected tmDCs with the session-averaged population activity in each head-directional bin (60 bins in total), following the method used in Vollan et al. [4]. Since tmDCs were mostly activated around the 180° phase in each theta cycle during straight runs [4] (Fig. S5), the time bin corresponding to this phase was used to extract the sweep angle in each theta cycle. Specifically, two inputs were fed into to the PV correlation decoder: first a *N* × *T* matrix (*M*^*PV*^) of temporally smoothed firing rates (Gaussian kernel with *σ* = 10ms), with *N* representing cell numbers and *T* representing the number of theta cycles; and second, an *N* × *P* matrix (reference matrix; *M*^*rPV*^) of session averaged angular tuning curves for head direction, with *P* representing the directional bins (*P* = 60). For the *t*_*th*_ theta cycle, the Pearson correlation coefficient was calculated between the *t*_*th*_ column of *M*^*PV*^ and each column of *M*^*rPV*^, resulting in a 1 × *P* vector representing the correlation coefficients with all angular bins (*T* × *P* correlation coefficients in total). The sweep angle in the *t*_*th*_ theta cycle was defined as the circular mean of *P* angular bin values, weighted by the *P* correlation coefficients matrix (dot product), hence producing a single scalar value representing the decoded sweep angle in each theta cycle.

We applied this process to each theta cycle resulting in a 1 × *T* vector representing all the decoded sweep angles in one recording session. These sweep angles are allocentric, being defined by room-referenced directional tuning curves. To get the egocentric directional sweep angle, we then subtracted the decoded sweep angle from the animal’s heading direction at the decoded time bin. Theta cycles for which the preceding egocentric internal direction pointed to the right of the head axis were grouped into the left-sweep class, whereas those whose preceding internal direction pointed to the left of the head axis were grouped into the right-sweep class. Kernel density estimation was then applied separately to the left-sweep and right-sweep groups to obtain their respective distributions of egocentric sweep angles (Fig. 2D). The peak of each distribution was extracted to represent the averaged sweep angles for left directional sweeps and right directional sweeps. Then, we took their absolute values and calculated their mean as the averaged directional sweep angle.

To calculate an overall averaged sweep angle in one recording session (irrespective of which module tmDCs were assigned to), we used the PV from all tmDCs in the PV correlation decoder (median=190 tmDCs in each session, IQR: [96-235]). To calculate an averaged sweep angle in each assigned module, we used the PV from each assigned module to decode their internal direction sweeps. Modules containing fewer than 20 tmDCs were excluded from analysis to prevent unreliable decoding. This resulted in 42 modules from 17 sessions in 10 animals with at least 2 modules involved in decoding (median= 82 tmDCs per assigned module, IQR: [55-110]).

Given that tmDCs within each assigned module fired at different phases of the theta cycle (Fig. S5), we first summed the spikes from all tmDCs in each module within each phase bin and identified the time window in which spike count was maximal. We then selected the decoded angles occurring within the same time window across all theta cycles and computed their mean. This yielded a sweep angle for each tmDC module, based on its preferred firing phase.

### Decoding locational theta sweeps in grid cells

The locational theta sweeps in grid cells were decoded using the same PV-correlation method as described above, with the PV from grid cell activities and the session-averaged spatial tuning maps as inputs to the decoder. The spatial tuning maps of grid cells were computed by binning positions into 2.5 × 2.5 cm spatial bins, calculating firing rates per bin, and smoothing the resulting map with a Gaussian kernel [4]. To calculate overall grid cell sweeps, we calculated the correlation between the PV of all recorded cells and the session-averaged spatial tuning maps (median= 741 cells per session, IQR: [606-992]). To calculate grid cell sweep in individual grid module, we conducted the same correlation but with the PV of grid cells in each grid module. Modules with less than 40 grid cells were excluded from the analysis to prevent unreliable decoding. This resulted in 48 modules from 19 sessions in 11 animals with more than 2 modules in decoding (median= 100 grid cells per module, IQR: [69-129]).

Within each theta cycle, the theta sweep trajectory was defined as the longest decoded trajectory in consecutive time bins, which was truncated to remove folds and maximize the Euclidian distance from the start point to the end point [4]. After calculating locational sweeps in every theta cycle, the session-averaged locational sweeps were computed by first interpolating each head-centred sweep trajectory at 50 time points linearly spaced from the beginning to the end of the trajectory. Interpolated sweeps were grouped into those that followed a right sweep or left sweep, before an average sweep was computed for each group by taking the median position at corresponding time points within the sweep [4]. The locational theta sweep angle was then calculated as the direction of the vector from the starting point to the endpoint of the session-averaged locational sweep, relative to the heading direction, and finally averaged across the left and right angles.

It is noteworthy that, although we observed increasing sweep angles across grid modules, the first grid module detected in each recording session was always designated as Module 1, even though it may not correspond to the anatomically most dorsal module. This does not violate the assumptions of the linear mixed-effects model, because our inference concerns within-animal module effects, and all statistical analyses were performed at the animal level. This is also the case in the analysis of increased internal-direction sweep angle from tmDCs.

### Analysing directional sweeps and locational sweeps across theta phase bins

To show that decoded internal direction sweeps continuously from side to side of heading direction across theta cycles, instead of flickering abruptly across the heading direction, as reported in Vollan et al. [4], we checked the decoded probability map as a function of theta phase. To do this, we binned each theta cycle into 20 phase bins (each bin is 18°) and binned the corresponding PV correlation matrix of directional sweep (described above) into 60 angular bins (each bin is 6°). Then we raised the correlation matrix to the 4^th^ power for better visualisation and normalised each column into a probability distribution. This resulted in a 20×60 probability matrices (PMs) of decoded direction in each theta cycle. We then averaged the PMs across all theta cycles in which the internal direction on the preceding theta cycle lay on the left side of the heading direction. We further averaged these mean PMs across all sessions, separately for modules 1, 2, and 3. This shows a right sweep starting from the heading direction at early theta phase (0°), propagating to the right side until middle theta phase (180°) and then sweeping backward to the heading direction until late theta phase (360°) (Fig. S3). This is also true by averaging PMs across all theta cycles in which the internal direction on the preceding theta cycle lay on the right side of the heading direction, which gives left sweeps.

To demonstrate that the locational sweep angle increases continuously within a theta cycle, we first selected locational sweep angles along the outward sweep trajectory. Only sweeps with 3–10 valid points and a path straightness score (computed for each time step of the recording by dividing net travel distance by cumulative travelled distance within a 2 s moving window; high values correspond to straight paths) greater than 0.5, obtained during running speeds above 20 cm/s, were included in further analysis. For the remaining sweeps, we interpolated each sweep trajectory into 10 points and calculated nine angular values of sweep direction, defined by the vectors from the starting sweep point to the second through tenth points, respectively. We then subtracted the heading direction to obtain nine egocentric sweep angles. These values were averaged across all selected sweeps within each recording session, separately for grid modules 1, 2, and 3. This yielded session-averaged locational sweep angles as a function of interpolated theta phase bins 1 to 9 (not exact phase bins, but close approximations). Finally, we averaged across all sessions from all animals, revealing a continuous increase in locational sweep angles across the theta cycle (Fig. S3 C).

Finally, to demonstrate that the directional sweep angle drives the continuous increase in locational sweep angle across a theta cycle, we first calculated the histogram of theta phases corresponding to the starting points of the selected locational sweeps and identified the first phase bin as the peak of this histogram. Starting from the subsequent phase bin to the first phase bin, we then calculated the directional sweep angles in the subsequent eight consecutive phase bins (18° per bin). These values were averaged across all theta cycles, separately for grid modules 1, 2, and 3. This procedure yielded session-averaged directional sweep angles as a function of theta phase bins that were approximately aligned with those used for the locational sweeps described above. Finally, we averaged across all sessions from all animals, revealing a continuous increase in directional sweep angles across the theta cycle (Fig. S3 D).

### Calculating theta skipping index in tmDCs

To quantify theta-skipping effect in tmDCs, we applied the method used in Brandon et al. (2013), fitting the autocorrelogram with a cosine wave of frequency *ω* and an interfering cosine wave of frequency *ω*/2:

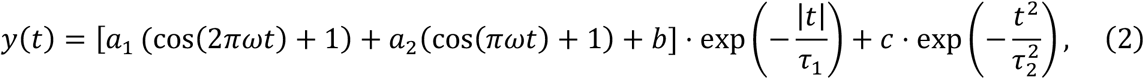

where *t* is the time lag of the autocorrelogram (in seconds), and *a*_1_, *a*_2_, *b, c, τ*_1_, *τ*_2_ are parameters determined by fitting. These parameters were constrained within the following ranges: a_1_ ∈ [0,100], a_2_ ∈ [0,100], *b* ∈ [0,100], *c* ∈ [0,0.8], ω ∈ [6,12], *τ*_1_ ∈ [0,8] and *τ*_2_ ∈ [0,0.05]. The interfering cosine wave was used to capture the theta-skipping effect on the autocorrelogram. The theta-skipping index (TSI) was then calculated as the difference between the first and second peaks on the fitted curve, normalized by the larger of the two:

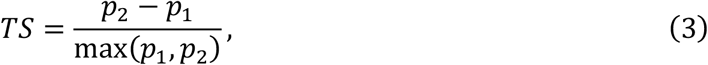

where *p*_1_ is the model value at one cycle with *t* = 2*π*/ω and *p*_2_ is the model value at two cycles with *t* = 4*π*/ω. This index is bounded between -1 and 1, with higher values indicating a greater degree of theta skipping. Note that TSI was calculated only based on spike trains during fast running periods with animals’ speed > 15 cm/s since stationary periods were not included in the published dataset.

### Computational modelling

The computational model consists of a two-layer continuous attractor network (CAN) in which tmDCs are modelled as a ring attractor and grid cells are modelled as a two-dimensional attractor on a neuronal sheet (GC attractor; see Ji et al. [16] for more details). The dynamics of the tmDC ring attractor are defined as:

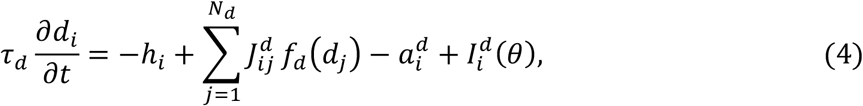

where *d*_*i*_ is the pre-synaptic input to *i*_*th*_ tmDC (100 simulated cells in total), *f*_*d*_(*d*_*j*_) is the firing rate of the j-th tmDC, 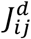 denotes Gaussian-like recurrent connections, 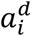 is the FRA term (defined below), and 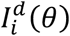 is the sensory input encoding the current head/movement direction:

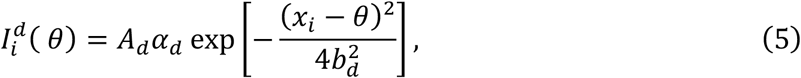

where *α*_*d*_ represents the theta modulation from the medial septum (see below) and *θ* is the current head/movement direction.

With firing rate adaptation, the activity bump sweeps from side to side around the current head/movement direction, with the sweep frequency controlled by the induced theta modulation.

The GC attractor is defined as:

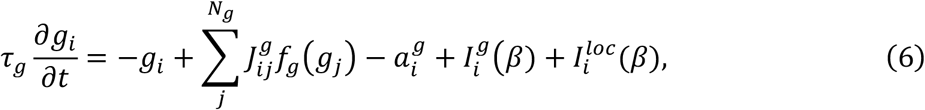

where *g*_*i*_ is the pre-synaptic input to the *i*_*th*_ grid cell (100 × 100 cells in total tiling the two-dimensional neuronal sheet), *f*_*g*_(*g*_*j*_) is the firing rate, 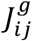 denotes the 2D Gaussian recurrent connectivity, 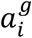 is the FRA term (defined below), *β* is location expressed in phase space (not physical space). Note that the GC attractor runs in a phase space instead of the physical space, so all the inputs has been converted from physical space to the phase space. 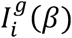 is the input from conjunctive grid cells, and 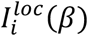 is the sensory input representing location information. 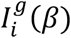 is expressed as:

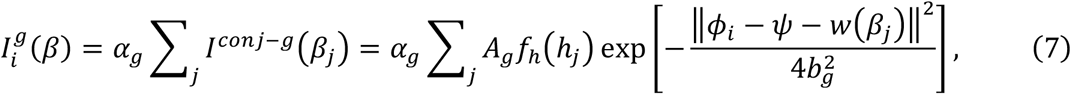

where *I*^*conj*−*g*^(*β*_*j*_) is the input from the *j*_*th*_ conjunctive-grid-cell group, *α*_*g*_ represents theta modulation (see below). Note that the *j*_*th*_ group of conjunctive grid cells will be activated when the *j*_*th*_ tmDC is activated in the ring attractor. The total input is a weighted sum of many 2D Gaussian bumps from all the conj-grid cells with the weights as the firing rate of upstream tmDCs *f*_*h*_(*h*_*j*_). *ψ* is the animal’s location in phase space. *w*(*β*_*j*_) is a vector with *w*(*β*_*j*_) = *w*_0_(cos *β*_*j*_, sin *β*_*j*_), where *w*_0_ represents the offset length, with the offset representing the grid phase offset between putatively connected conjunctive grid to pure grid cells, as shown in Vollan et al [4]. Eq. (7) means activation of conjunctive grid cells corresponding to direction *β*_*j*_ generates a shifted Gaussian input centered around *ψ* + *w*(*β*_*j*_). The location-dependent input 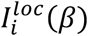 is expressed as:

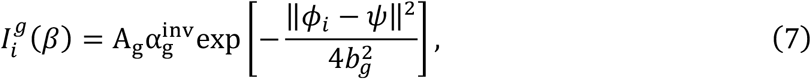

where 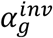 is theta modulation with an opposite phase compared to *α*, mimicking encoding and retrieval occurring at opposite phases of the theta cycle [34]. Thus, in early theta phases, the GC attractor is pulled toward the current location *ψ*, while in late phases it is driven toward the offset input from conjunctive grid cells, generating directional sweeps along the internal direction.

Theta modulation of tmDC is:

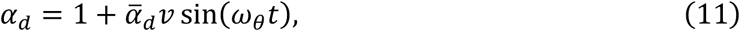

where the oscillation magnitude scales linearly with the angular speed *v*, with 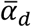 a scaling factor.

The theta modulation on the GC attractor is:

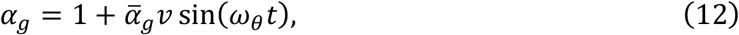

where the oscillation magnitude also scales linearly with the running speed *v*, with 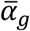 a scaling factor.

Firing rate adaptation has the following dynamics:

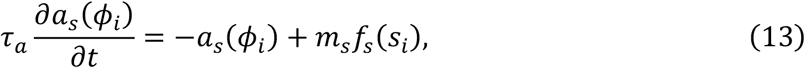

where *a*_*s*_(*ϕ*_*i*_) represents FRA in the *i*_*th*_ cell with *s* ∈ {*d, g*}. *d* represents tmDCs and *g* represents grid cells. *f*_*s*_(*s*_*i*_) represents the firing rate of the *i*_*th*_ cell. *τ*_*a*_ is the time constant of FRA, which is much larger than *τ*_*d*_ and *τ*_*g*_, indicating slow dynamics. *m*_*s*_ is FRA strength, which can differ between tmDCs and grid cells. We systematically varied *m*_*s*_ to model the dorsoventral increase in FRA observed empirically [26].

For the parameter settings, the simulated model consists of 100 tmDCs (*N*_*d*_) and 10000 grid cells (*N*_*g*_) arranged on a 2D neuron sheet. Time constants for both tmDCs (*τ*_*d*_) and grid cells (*τ*_*g*_) are set to 10 ms, as is typical in rate-based models. For tmDCs, recurrent connections are modelled as Gaussian profiles with a standard deviation (*b*_*h*_) of 0.4 radians. For grid cells, recurrent connections are modelled as 2D Gaussian profiles with a standard deviation (*b*_*g*_) of 0.8, corresponding to approximately13% of the width of the 2D neuronal sheet. The offset value projected from conjunctive grid cells to pure grid cells, *w*_0_, is set to 1/9, corresponding to roughly1.8% of the width of the sheet. The time constant for firing rate adaptation, *τ*_*a*_, is set to a much larger value of 100 ms compared with the time constant *τ*_*g*_ and *τ*_*d*_. This means it takes approximately 100 ms for a neuron’s firing rate to adapt (i.e., decrease) in response to a sustained stimulus.

To model the increase in sweep angles along the dorsoventral axis (where adaptation strength increases [26]), the adaptation strength of tmDCs (*m*_*d*_) varied across [10,12,14,16,18, 20], with 10 corresponding to the most dorsal part and 20 to the most ventral part of the mEC/parasubiculum.

### Statistical Analysis

All statistical analyses were performed in Python (version 3.12). Statistical analyses in Fig. 1-3 and S1&4 were performed using **statsmodels** package (MixedLM implementation). Statistical significance was defined as α = 0.05, and all tests were two-sided. Exact P values, test statistics, and sample sizes are reported in the main text or corresponding figure captions. To account for repeated measurements across animals, linear mixed-effects models were fitted with sweep angle as the dependent variable and module identity as a categorical fixed effect. Animal identity was included as a random intercept to account for clustering of observations within animals. Models were fitted using maximum likelihood estimation when comparing nested models and restricted maximum likelihood otherwise. The overall effect of module was assessed by comparing the full model to a null model containing only the intercept and random effect, using a likelihood ratio test. The test statistic was computed as twice the difference in log-likelihoods and evaluated against a χ^2^ distribution with degrees of freedom equal to the difference in the number of fixed-effect parameters. Pairwise comparisons between modules were obtained by refitting the model with alternative reference levels, and regression coefficients (β), χ^2^ statistics, and exact P values are reported.

Pearson correlation in Fig. 4B coefficients were computed using the **scipy.stats** module in Python. Correlation strength (r) and corresponding two-tailed P values were calculated to assess linear associations between variables. Statistical significance was defined as α = 0.05. Sample size, correlation coefficients, and exact P values are reported in the main text or figure legends.

## Acknowledgments

We thank the Moser lab for making the experimental data from Vollan et al. (2025) available online. We thank Motoharu Yoshida and Michael E. Hasselmo for providing the patch-clamp data. We thank Callum Marshall, Tianhao Chu for discussion about the computational model. We thank Daniel Bush and Kate Jeffery for valuable comments.

## Author contribution

Conceptualization: ZJ, NB

Data analyses: HZ, ZJ, RS

Computational modelling: ZJ Supervision: ZJ, NB

Writing—original draft: ZJ, HZ

Writing—review & editing: ZJ, NB

### Data and materials availability

All data are available in the main text or the supplementary materials. They are publicly available at: https://doi.org/10.25493/R5FR-EDG. Code for reproducing all the results in the main text will be available upon publication.

## Supplementary Figures

**Fig. S1:**
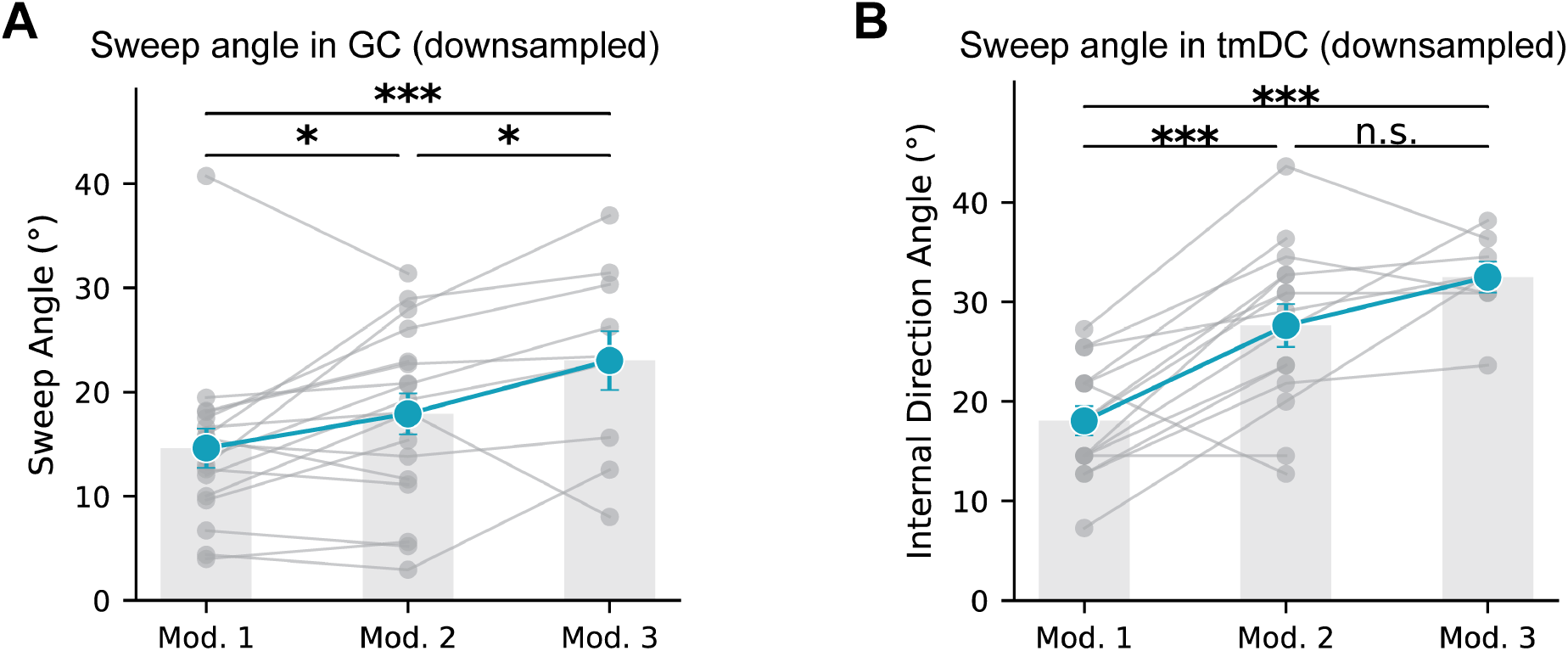
Topography of sweep angles in grid cells and tmDCs along the mEC/parasubicular DV axis after sub-sampling the number of cells in each module to match the number of cells in decoding. **(A)** Mean sweep angle in grid cells across three grid modules. Each dot represents the sweep angle decoded from one module (19 sessions from 11 animals). Grey lines link the mean sweep angles from assigned modules in the same recording session. A linear mixed-effects model yielded *χ*^2^(2) = 14.7, p = 6.4 × 10^−4^. Averaged sweep angles were module 1, 14.6°; module 2, 17.9°; module 3: 23.0°. Module 2 vs module 1: β = 3.32, p = 0.03. Module 3 vs module 2: β = 4.75, p = 0.015. **(B)** Mean sweep angle in tmDCs across three assigned modules. Each dot represents the sweep angle decoded from one assigned module (17 sessions from 10 animals). Grey lines link the mean sweep angles from assigned modules in the same recording session. A linear mixed-effects model yielded *χ*^2^(2) = 27.5, p = 1.1 × 10^−6^. Averaged sweep angles were module 1, 18.1°; module 2, 27.6°; module 3: 32.5°. Module 2 vs module 1: β = 9.58, p = 3.5 × 10^−7^. Module 3 vs module 2: β = 4.15, p = 0.079).

**Fig. S2:**
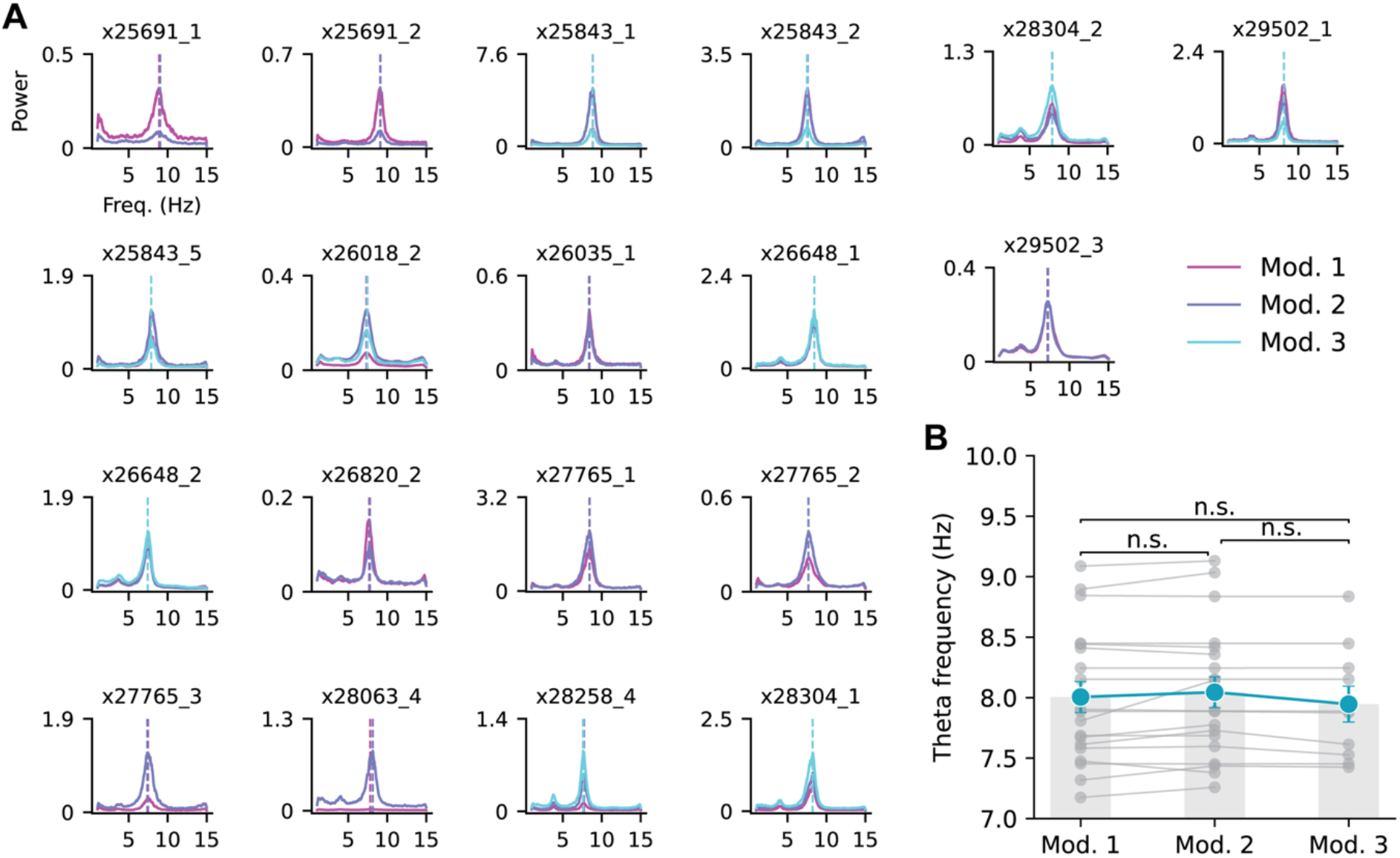
Theta frequency did not show a gradient along the mEC dorsoventral axis. **(A)** Power spectra calculated from summed spikes within each grid module. Each panel represents a recording session from one animal, with the animal and session ID indicated in the panel. Dashed lines mark the peak theta frequency. **(B)** Peak theta oscillation frequency across three modules. Each dot represents the peak frequency derived from the summed spike activity with a grid module (19 sessions from 10 animals). Grey lines link the mean peak theta frequency of grid modules detected within the same recording session. A linear mixed-effects model yielded *χ*^2^(2) = 4.77, p = 0.09. The mean peak theta frequencies were 8.0 Hz for module 1; 8.0 Hz for module 2; 7.9 Hz for module 3.

**Fig. S3:**
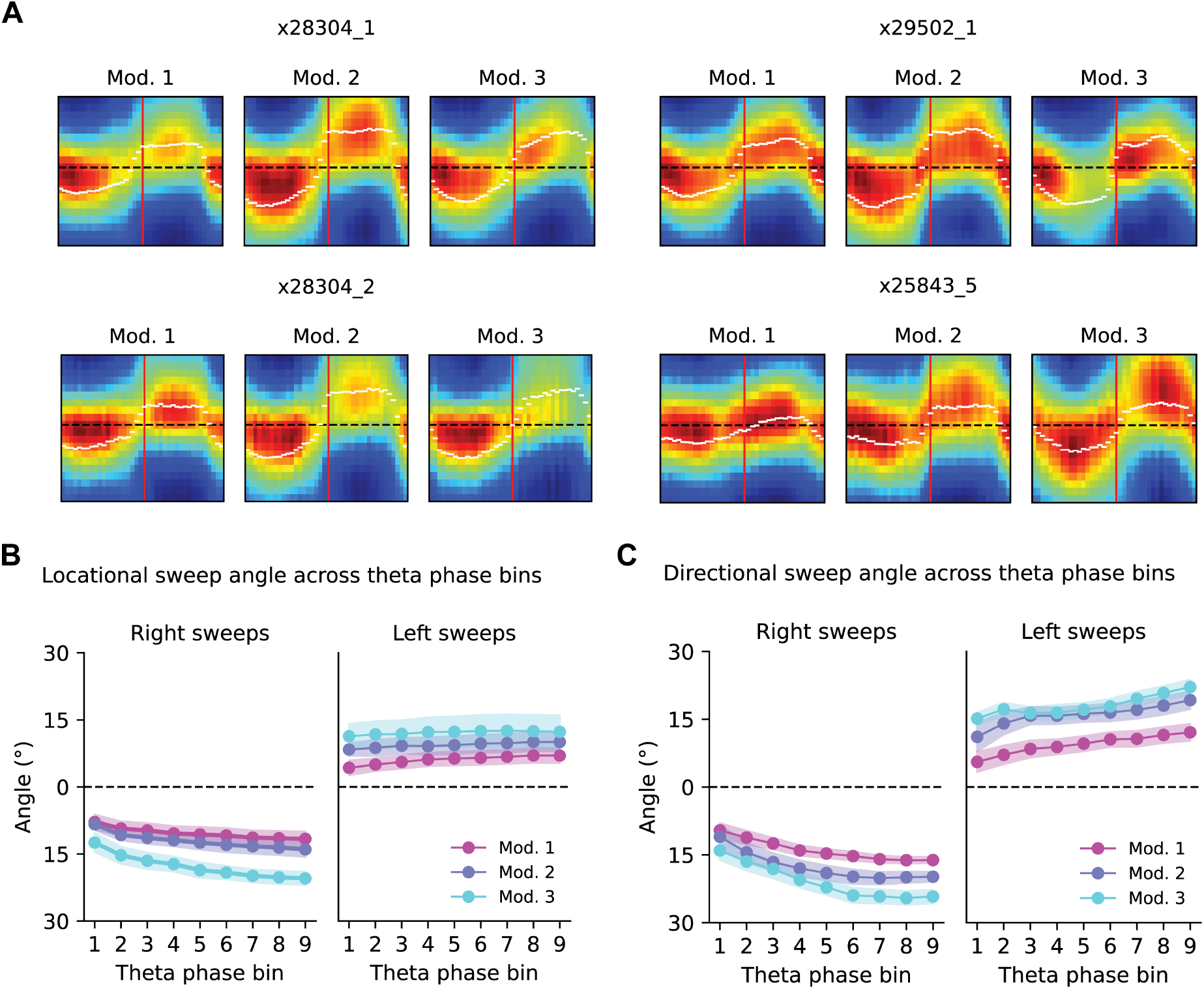
Internal directional sweeps evolve continuously, rather than flicker around the heading direction, and drive continuous locational sweeps. **(A)** Four examples of directional sweeps in individual recording sessions. The red line separates theta cycles, with the left portion representing cycles corresponding to right sweeps and the right portion representing cycles corresponding to left sweeps. The dark dashed line marks the heading direction (reoriented to zero), and the white dashed line marks the mean directional sweep angle at each theta phase. **(B)** Session-averaged locational sweep angle as a function of theta phase bin (showing roughly only half of the theta cycle, since backward sweeps were cut off). Colours from magenta to blue indicate the locational sweep angle in grid modules 1 to 3. The left panel shows rightward sweeps, whereas the right panel shows leftward sweeps. The first phase bin corresponds to the sweep angle calculated from the first two points of the sweep trajectory, and the last phase bin corresponds to the sweep angle calculated from the start and end points of the sweep trajectory (see Methods). Backward sweeps were excluded from all calculations. The shaded area indicates the s.e.m. of locational sweep angles in each module, with 23 sessions in module 1, 19 sessions in module 2, and 10 sessions in module 3. **(C)** Session-averaged directional sweep angle as a function of theta phase bin. Colours from magenta to blue indicate the directional sweep angle in assigned modules 1 to 3. The shaded area indicates the s.e.m. of sweep angles in each module, with 19 sessions in assigned module 1, 20 sessions in assigned module 2, and 10 sessions in assigned module 3.

**Fig. S4:**
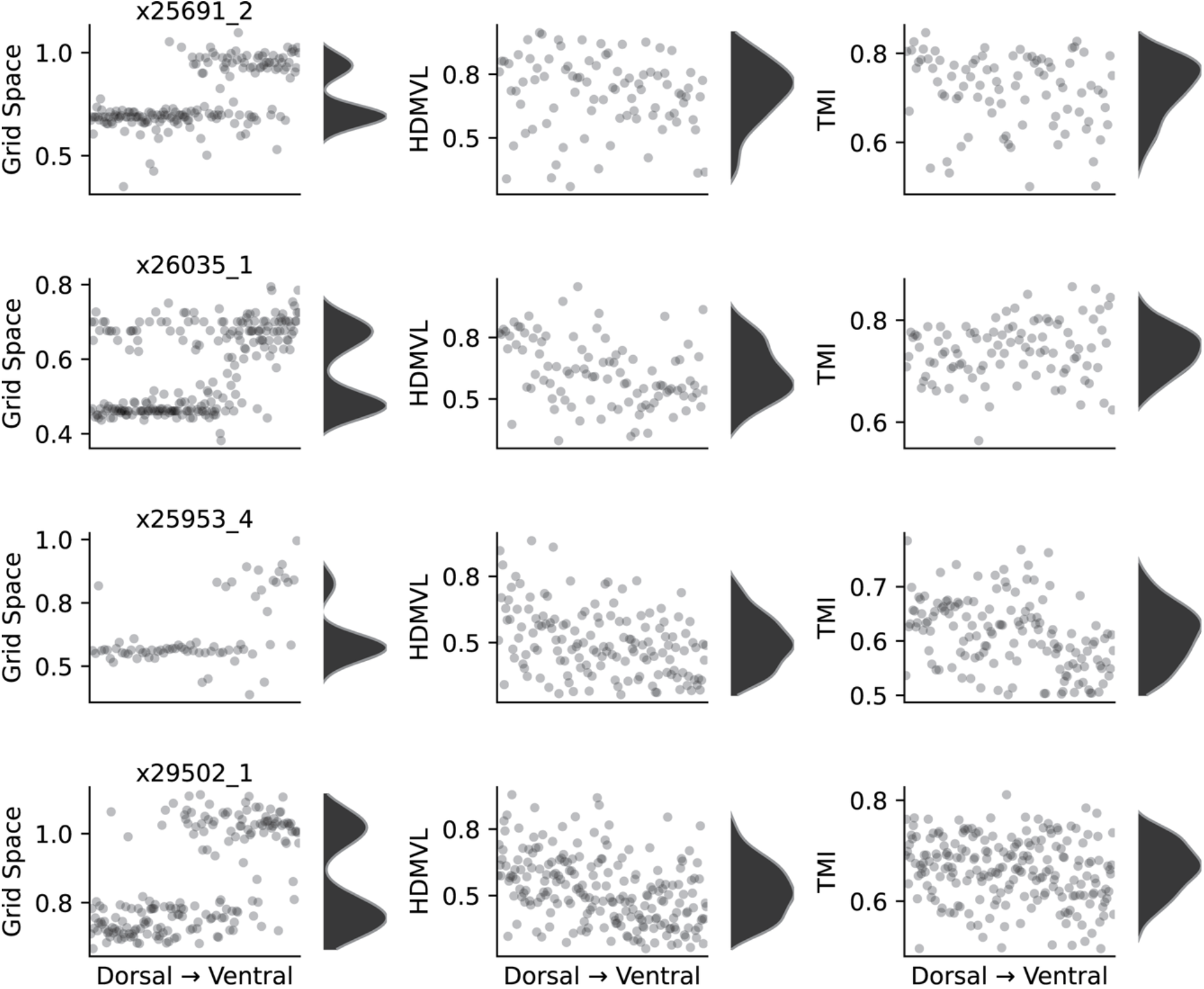
Gradient of single-cell firing features along the mEC/parasubicular dorsoventral axis from four animals. Each dot is one cell, and each row is one recording session. Column 1: distribution of grid spacing at different recording sites along the dorsoventral axis. Column 2: distribution of head-direction mean vector length (MVL) calculated for each tmDC. Column 3: distribution of the theta modulation index calculated for each tmDC.

**Fig. S5:**
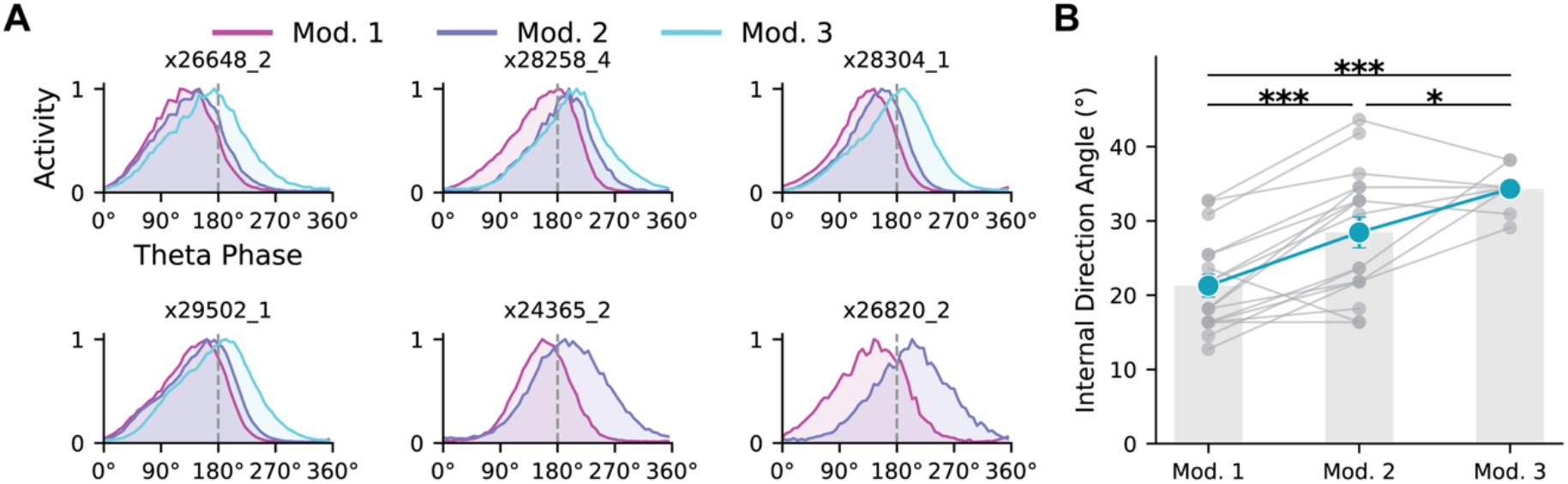
Topography of sweep angles in tmDCs after correcting for their preferred firing phase relative to theta. **(A)** Examples of preferred firing phase of tmDCs across assigned modules from six animals. Theta phase was extracted from the population spiking activity of all units recorded in the session, with 0°/360° corresponding to the minimum firing. Activity was computed by summing spikes from tmDCs (within each assigned module) in each theta-phase bin and normalising by the maximum spike count across bins. Dashed lines indicate 180° for reference. **(B)** Mean sweep angle across the three assigned modules. Each dot represents the sweep angle decoded from one module (17 sessions from 10 animals). Grey lines link the mean sweep angles from assigned modules in the same recording session. A linear mixed-effects model yielded *χ*^2^(2) = 25.3, p =3.2 × 10^−6^. Averaged sweep angles were module 1, 21.3°; module 2, 28.4°; module 3: 34.3°. Module 2 vs module 1: β = 7.16, p = 7.9 × 10^−6^. Module 3 vs module 2: β = 4.39, p = 0.041.

